# AraMultiOmics: a tool for generating multi-omics features for downstream statistical analysis to infer the molecular basis of symbiosis among non-host plant Arabidopsis thaliana, host plant, and arbuscular mycorrhizal fungi

**DOI:** 10.1101/2023.11.01.565228

**Authors:** Jee Eun Kang

## Abstract

Arbuscular mycorrhizal fungi (AMF) are symbiotic microorganisms that colonize plant roots, promoting plant growth and improving soil quality. A large number of studies have focused on investigating the communication between host-plants and AMF. Recent investigation in molecular evolution related to arbuscular mycorrhizal symbiosis in Arabidopsis thaliana (A. thaliana) have enabled scientists to perform comparative multi-omics analyses between A. thaliana and host-plants. Although there is a vast amount of omics data available for A. thaliana, most AM-related information comes from differentially expressed genes (DEG) identified in transcriptome studies. To address this gap, we developed AraMultiOmics, a useful tool for integrative analysis of multi-omics data of A. thaliana. It consists of 10 modules: 1) epigenetic regulations in protein–nucleic acid interaction (PNI), 2) DNA structure and metal binding profile, 3) transcription factor (TF) binding profiles, 4) protein domain–domain interaction (DDI), 5) profiling of interactions of protein-metal and of protein-ligand with complex structures (PLP) based on alignment of similar protein structures, 6) carbohydrate-lipid-protein interaction (CLP)- analysis of lipidome-proteome-glycoscience, 7) metabolic pathway analysis, 8) multiple omics association study, 9) GO/PO analysis, and 10) Medicago COG information. These analyses are conducted in comparison with the COG of Medicago truncatula (M. truncatula). To facilitate the inference of AM-driven changes and of AM derived molecules during AM symbiosis, the program provides a convenient means to generate datasets with important features that can be conjoined with various downstream statistical methods. We have included demonstrations on how to create comparative datasets, and the program codes are freely available for download at www.artfoundation.kr.

## Introduction

Arbuscular mycorrhizal (AM) symbiosis is one of the oldest plant-fungal mutualisms that promote plant growth and improve soil quality [1,2]. In the last decade, significant progress has been made in understanding molecular mechanisms underlying communication between host-plants and arbuscular mycorrhizal fungi (AMF). However, the non-host plant’s interaction with other members of the network remains largely unknown. Recent studies have indicated that the presumed non-host plants can develop rudimentary AM phenotypes [3]. Although non-host plant A. thaliana does not form a typical AM structure, AM colonization brings major changes in transcriptome [4]. It is speculated that A. thaliana might have lost the AM trait throughout evolution [5]. A. thaliana, modified with the key AM TF-IPD3, exhibited the expression of AM-related gene networks, suggesting the retention of the molecular machinery for symbiosis in A. thaliana. Additionally, it has been implied that CERK1 with LYMs and PEN1/VAMP721 may play a role in the perception of Nod factor analogs and vacuolar components of the SNARE complex essential for arbuscule accommodation, respectively [6-8]. Strigolactone biosynthesis genes belonging to early signaling between A. thaliana and AMF were activated in A. thaliana that was co-planted with M. truncatula within the mycorrhizal network [4].

Members in the AM network exchange fungal diffusible factors, root exudates, and extracellular vesicles [9]. Recent studies indicated that AM-related molecules caused significant changes to the overall transcriptome. However, their physiochemical properties remain elusive. Despite the vast amount of omics data generated for A. thaliana, information related to AM symbiosis is predominantly derived from DEGs identified in transcriptome studies. A large proportion of DEGs from plants during AM symbiosis belonged to cell communication, transporters, proteins involved in lipid and amino acid metabolism, small-secreted peptides, and TFs [4]. We need to infer signature patterns of molecular interactions during AM symbiosis based on existing data. For instance, properties of AM-derived molecules may be inferred from comparative analyses among host-plants and non-host plants. For example, this involves examining changes in PNI and protein-protein interaction (PPI) due to increased accessibility of transition metals and to availability of new signaling molecules with complex structures, analyzing glycosylation/glycan structure and the gene expression of enzymes involved in cell wall and lipid metabolism, and assessing membrane lipid composition/thickness/ tilt angles of membrane proteins engaged in transporting AM-derived molecules [10-13]. We developed the AraMultiOmics that consisted of 10 modules where each module contains features derived from existing databases and software: 1) protein binding sites/open chromatin regions identified from chip-seq and their derivatives (ChIP-Hub, JASPAR, PCSD (Plant Chromatin State Database)), 2) DNA structure and metal binding profile (cran R gquad package (v2.1-2), QGRS program, and mebipred program), 3) DNA-TF binding profiles (TF-COMB (TF Co-Occurrence using Market Basket analysis. V0.1.0)), 4) Protein DDI and membrane protein orientation (3did (three-dimensional interacting domains) and OPM (Orientations of Proteins in Membranes)), 5) PLP with protein structure similarity search (Alphafold database, foldseek2, PDB, PDBBind), 6) CLP (supplementary data from Bahammou et al., Radrich et al., and Burks et al., Glycosmos, dbCAN3, HMMER package, CSDB-GT, Amigo GO), 7) metabolic pathway analysis (PMN (Plant Metabolic Network)), 8) DNA methylation and gene expression associated with ion concentration phenotype (Epigenome-wide association study (EWAS), transcriptome-wide association study (TWAS), and muti-omics association (AtMAD), 9) GO, PO, IPD3 DEG type classification (TAIR, Hornstein et al.), 10) Medicago COG (String database) [5,13-42]. To compare the effects of AM molecules on PNI and PPI between host and non-host plants, we incorporated existing programs for predicting DNA structures. These structures induce conformational changes cooperatively with metals, and we also incorporated tools for assessing metal binding profiles of proteins [19,20]. To make inferences on the properties of AM-driven molecules including those with complex structures, we implemented PLP module based on Alphafold database, foldseek2, PDB, and PDBBind, and CLP module based on lipidome-proteom-glycoscience-mRNA-seq data, specifically frequencies of lipid and protein modifications, glycoconjugate structure information, glycosyltransferase reactions, metabolic networks from Aracyc and KEGG databases, GO/pathway enrichment analysis within each gene cluster/module based on gene co-expression data from mRNA-seq datasets, and carbohydrate active enzyme/substrate prediction. To infer the properties of membrane proteins whose structural changes influence signal transduction, we included information on membrane protein orientation in lipid bilayer from the OPM database [13]. To facilitate multi-module analyses (e.g., transcription-related PNI modulated by metal-driven changes in DNA structures and protein interaction, sequence diversity of metal binding in DDI, metal-driven protein oligomerization/crystallization, and correlations between metal homeostasis and gene expression of membrane protein), we unified relational keys of all modules by gene ID/domain ID [34-38]. Sequence information was included for further analysis.

To demonstrate how to create comparative datasets for AM symbioses from A. thaliana and M. truncatula, we included three simple analyses. In the first one, we selected COGs (e.g., early nodulin-like protein, F-box protein, probable WRKY transcription factor, Auxin transporter, ATPase) which contained at least one metal in their COG description. We classified COGs based on DEG types (none, up/down-regulation, both) during AM symbiosis from M. truncatula, and retrieved genes belonging to the same COGs from A. thaliana with IPD3 restoration. We merged various features (e.g., DNA structure, protein’s metal binding profiles, OCR information of various experiments, OCR-tissue maps, PCSD) from modules and combined them into datasets In the second one, we selected enzymes in volatile organic compound reactions (VOC), histidine biosynthesis genes (AtHISN1B, AtHISN2, AtHISN1A, AtHISN6B, AtHISN3, AtHISN5A, AtHISN5B, AtHISN4, AtHISN6A, At-HISH, AtHISN8), and homologous genes (CYP76C1, CYP76C2, CYP76C3, CYP76C4, CYP76C5, CYP76C6, GUS1, GUS2, GUS3, CSLG1, CSLG2, CSLG3, VTE1, VTE5, HAI1, HAI2, HAI3). We generated datasets with various modules-PNI, DDI, PLP, and metabolic pathway analysis. In the third one, we created datasets according to GO/PO categories (e.g., signal transduction, subcellular localization sorting, defense and cell rescue, and abiotic stimuli and development). Then, we included examples of downstream statistical analyses-clustering and classification methods for demonstration purposes [4,5]. The program provides a convenient means to generate datasets with various features that can be conjoined with downstream statistical analyses to elucidate the molecular interactions of AM symbiosis. The program codes and datasets of the AraMultiOmics are available for download at www.artfoundation.kr.

## Methods

AraMultiOmics consists of 9 modules created based on existing databases and programs.

### 2.1. Protein-DNA interaction module

Information contained in this module mainly focused on chromatin-related epigenetics and TF binding properties. The module included protein binding sites/open chromatin regions from more than 470 experimental datasets, mainly ChIP-seq and DNase-seq and their derivatives provided by the database-ChIP-Hub [14]. ChIP-Hub provided tissue specificity information on SRA experiments (e.g., various histone marks) and scores for enhancer regions. We re-organized open chromatin regions (OCR) data according to type (promoter/enhancer), and gene regions (e.g., intron, UTRs) per gene. We mapped genes with approximately 450 OCR SRA accessions and tissues (flower, inflorescences, leaf, root, root hair, root non-hair, seed, seed coat, seedling, and shoot). The bedtools-intersect function was used to identify the JASPAR motif regions in OCR data [15]. We mapped JASPAR motifs with TF domains according to the A. thaliana file provided by JASPAR. We used cran R TFBSTools to calculate similarity scores between JASPAR2014 motif sets and entries in the Place database [17,18]. Additionally, we incorporated information from PCSD (Plant chromatin state database) that provided chromatin states per gene calculated based on numerous chip-seq and atac-seq data [16]. We incorporated atac-seq data of phloem and leaf from Tian et al. and calculated the sequence length of accessible chromatin regions per gene [33].

In addition, genome-wide DNA methylation data of the AraPheno study 16 (Ion concentration-arsenic, cadmium, cobalt, copper, potassium, magnesium, manganese, molybdenum, Nickel, phosphorus, rubidium, selenium, sulfur, zinc) from the AtMAD database (EWAS) were incorporated with distinction of type and gene region [32-34]. Genes in GO analysis were mapped to various terms (e.g., xylem and phloem pattern formation, transmembrane transporter activity) [47,48].

### 2.2. DNA structure and metal binding profile module

DNA structures (G quadruplex, H-DNA, slipped motifs, short tandem repeats, triplex-forming oligonucleotides, and Z-DNA) of genes from A. thaliana and M. truncatula were predicted with cran R package-gquad and QGRS program [19]. Metal binding profiles (Ca, Co, Cu, Fe, K, Mg, Mn, Na, Ni, Zn, and metal binding potential) of proteins were predicted with the mebipred program [20]. We calculated the frequencies of distances from the start of the gene to DNA structure, considering ranges such as 1-99, 100-199, 200-299, 300-399, 400-499, 500-599, 600-699, 700-799, 800-899, 900-999, 1000-1499, 1500-1999, and greater than 2000. The module included parsing and mapping functions to integrate the results from the aforementioned programs.

### 2.3. DNA-TF binding module

The Cran R package, TF-COMB, was employed to identify co-occurring TFs based on chip-seq, atac-seq, and their derivatives [21]. TF-COMB predicted co-occurring TF pairs, distances between TFs, orientations of stranded regions, and differential co-occurrences under distinct conditions (github:loosolab/TF-COMB). For chip-seq input, we selected the SRP045296 project-a comprehensive atlas of Arabidopsis regulatory DNA (DAP-seq). Additionally, for atac-seq input, we used the SRP300093 project (atac-seq) focused on the evolution of conserved noncoding sequences. The TF binding site map on a genome-wide scale was obtained from ChIP-Hub. The program was executed in R, and the results were integrated into the module.

### 2.4. Protein domain-domain interaction module

We integrated features derived from the 3did database into the module. 3did collected 3D protein structures in PDB and clustered domain-domain interfaces in PPI into interaction topologies [22]. 3did provided information on residue membership belonging to each topology group and on fractions of how many binding partners used each residue. The module included functions to retrieve a sum of occurrences of each amino acid in topologies. We categorized a DDI group with the same PFAM architecture (e.g., Dom1-Dom2-Dom3-DPep1-DPep2) into two levels. Firstly, we categorized a DDI group according to the number of different domains in the PFAM architecture: only one PFAM domain (e.g., Dom1), multiple PFAM domains (e.g., Dom1-Dom2), and both (e.g., Dom1 and Dom1-Dom2). Secondly, we sub-categorized them based on whether they included DDIs with redundant domains (e.g., Dom1-Dom2-Dom1-Dom3-Dom2) or DMI with redundant peptides (DPep1-DPep1-DPep2). The module contained functions to map domains with HMM profiles in different formats provided by PFAM. We incorporated metal binding profiles (Ca, Co, Cu, Fe, K, Mg, Mn, Na, Ni, Zn) per protein domain from the mebipred database [20]. We incorporated information on protein orientation within the lipid bilayer, including tilt angles and membrane thickness, for membrane proteins obtained from the OPM database [13]. In addition, we incorporated PPI data from the string database.

### 2.5. Protein–ligand profile (PLP) module based on protein structure similarity search

Protein structure information of A. thaliana was provided by Alphafold database [23]. We used foldseek2 program to create searchable PDB and PDBBind databases, and to find regions of local similarity between structures of Alphafold and the created database [24]. The module included functions to retrieve protein structure information (e.g., quaternary structure, geometry and sterochemistry of atoms in close contact, rigid-bond restraints, sphericity, non-bonded contacts) and protein-ligand information (e.g, important sites of protein-protein residue in contact with ligands), protein secondary structures (helix, sheet, those in-between), and details of ligand properties (e.g., frequencies of elements-Al:As:Br:Ca:Cd:Cl:Co:Cs:Cu:Du:Fe:Hg:Li:Mg:Mn:Na:Ni:Pb:Se:Sr:Yb:Zn:B:C:F:H:I:K:N:O:P:S:U :Y, frequencies of bond-single:double:triple:amide:aromatic, six properties of atom-aromatic:leaving atom:right/left hand of stereocenters:flags for backbone:n_terminal:carboxyl-terminus:amino-terminus, frequencies of atom properties in SMILES string-types of bonds:branching:special case atoms:stereochemistry). We mapped PDB entries per Uniprot IDs and combined information of protein secondary structures and ligands. Based on foldseek2 output, protein-ligand information, protein secondary structures, ligand details, and Uniprot combined information can be retrieved per residue in aligned portions between Alphafold structure of A. thaliana and target PDB.

### 2.6. carbohydrate-lipid-protein interaction (CLP)

Bahammou et al. provided information on plasma membrane lipidome and proteome, specifically normalized spectral abundance factor values from the six fractions used in protein identification, localization, N-myristoylation, S-palmitoylation, prenylation (S-farnesylation and/or S-geranylation), Glycosylphosphatidylinositol (GPI) anchor prediction, and GRAVY. [27]. Glycosmos provided annotation on proteins, specifically structures of glycans-carbohydrate structures attached to proteins and lipids and glycoconjugates, frequencies of N-glycosylations, and GlyTouCan IDs [28]. dbCan developed program to predict substrate structures of carbohydrate-active enzymes with HMMER package [29, 30]. It mapped enzyme subfamilies with CAZY ID and E.C. number.

CSDB-GT (Carbohydrate Structure Glycosyltransferase Database) provided informatin on enzyme belonging to glycosyltransferases, CAZY family, gene cluster, structures of substrate and product, synthesized bond, and donor structures [41]. Radrich et al. provided information on compound and reactions of metabolic networks mapped to both Aracyc and KEGG databases [39]. Burks et al. provided GO and pathway enrichment analysis data within each gene cluster/module of correlated gene expression based on more than 20000 mRNA-seq datasets [50]. Amigo Gene Ontology data were incorporated to annotate GO and pathway analysis [42]. We developed the CLP module by integrating the above information to provide users with carbohydrate, lipid, and protein interaction data which can be retrieved with gene name.

### 2.7. Metabolic network analysis module

The module contained plant metabolic pathways-gene, protein, reaction, compound, and pathway provided by PMN (Plant Metabolic Network) databases [27]. Feature variables of compounds were created based on three different fields named “TYPES”, “CHEMICAL-FORMULA”, and “STRUCTURE-GROUPS”. The variable, created based on a chemical formula, had presence/absence values of the following elements: As, Br, C, Ca, Cd, Cl, Co, Cu, Fe, H, I, K, Mg, Mo, N, Na, Ni, O, P, S, Se, Zn. We mapped PMN compound ID with KEGG compound ID and created additional features, such as frequencies of a compound in 438 reaction groups and frequencies of “TYPES” of PMN compounds in 438 reaction groups, based on reaction classification with KEGG atom types from the RCLASS database [49]. To facilitate integration with other modules, reaction data were re-organized into different data structures (reaction type, pathway, substrate groups, product groups) with reaction enzyme as key.

### 2.8. Muti-omics association module

The module contained multi-omics association information of A. thaliana provided by AtMAD [32]. In EWAS (epigenome-wide association study) data, DNA methylation data were associated with phenotypes. We selected the phenotype AraPheno Study16 (Ion concentration), in which A. thaliana accessions were grown in a controlled environment with the following elements added at subtoxic concentration: arsenic (As75), cobalt (Co59), molybdenum (Mo98), selenium (Se82), zinc (Zn66), nickel (Ni60), copper (Cu65), magnesium (Mg25), phosphorus (P31), rubidium (Rb85), cadmium (Cd111), manganese (Mn55), potassium (K39), sulfur (S34) [-30]. We re-organized EWAS data according to type, gene regions, overlapping regions of OCR peak sites, metal types, and methylation types-CG, CHG, and CHH (H=A,C,T). Gene expression data of the same phenotype (Study16: ion concentrations) were extracted from a transcriptome-wide association study (TWAS). In addition, we incorporated phenotype-pathway association data for the same phenotype.

### 2.9. GO, PO, IPD3 DEG module

GO and PO analysis data from the TAIR database were incorporated [35]. We selected 17 categories from GO: abiotic, RNA, biotic, secretion, cytoskeleton, signal, Defense, skeleton, hormone, subcellular localization sorting, membrane_transport, transport, metabolism, vesicular, phloem, and xylem, polarity. We selected 10 categories from PO: defense, metabolism, phloem, subcellular localization sorting, RNA, secretion, signal, transport, vesicular, and xylem. In addition, we incorporated IPD3 DEG module data, where module classification was obtained by applying weighted gene co-expression network analysis to transcriptome data of A. thaliana samples [5].

### 2.10. Medicago COG module

The Clusters of Orthologous Groups of proteins (COG) database from the String database was used to map gene IDs to COG IDs and their corresponding descriptions for both M. truncatula and A. thaliana [36]. Intersected COGs belonging to both species were retrieved. COGs were categorized into four types based on whether they contained DEGs from M. truncatula of AM symbiosis: none, up-regulation, down-regulation, and both up- and down-regulations. IPD3 DEG module types of genes of A. thaliana belonging to the intersected COGs were provided.

### 2.11. Demonstration of Program Usage

#### 2.11.1. Comparative dataset generation based on DEGs of AM symbiosis from M. truncatula and A. thaliana

We selected four metals: sodium, potassium, nickel, and manganese, and retrieved COGs whose descriptions contained at least one of the metals. We categorized COGs from M. truncatula based on whether they contained only up-regulated, down-regulated, both, or no DEGs. From A. thaliana with IPD3 restoration, we retrieved DEG module types of genes belonging to the same COGs. We retrieved DNA structures, protein metal binding profiles, OCR SRA experiment IDs, OCR tissue information, and PCSD data and combined them into datasets.

#### 2.11.2. Analysis of DEG genes from A. thaliana with transgene knock-in mutant of IPD3 from Hornstein et al

Among DEGs from A. thaliana with IPD3 restoration, we selected the homologous genes-CYP76C1, CYP76C2, CYP76C3, CYP76C4, CYP76C5, CYP76C6, CSLG1, CSLG2, CSLG3, VTE1, VTE5, HAI1, HAI2, and HAI3. With the selected genes, we retrieved PCSD chromatin states and applied clustering methods such as FarthestFirst.

To compare the metabolic pathways of VOC, we selected genes in the glycolysis pathway and 4 different metabolic pathways related to the VOC-shikimate pathway, mevalonate (MVA) pathway, lipoxygenase (LOX) pathway, and pathway for linoleic acid. We also included other PAL homologous genes (PAL1, PAL2, PAL3) [50]. We retrieved PCSD chromatin states and OCR SRA accession-tissue data, substrate/product compound variables, DDI category data (redundant and non-redundant DDI/DMI), and max topology number of domain dimers. We applied the clustering methods-“cobweb” and “canopy” in the Weka program [51-54]. Default value parameter set was used with cobweb algorithm: acuity:1.0; cutoff:0.0028; random number seed:42. Default value parameter set was used with canopy algorithm: number of clusters:2; max-candidates:100; how often to prune low density canopies:10000; minimum canopy density:2; T2 distance:-1.0; T1 distance:-1.5; random seed number:1. Frequencies of DNA structures and protein metal binding potentials of PFK genes from two species were compared.

Available Alphafold structures of the homologous genes were HAI2, CYP76C3, CYP76C4, CYP76C5, CYP76C6, PAL2,VTE7. We used foldseek2 program to create the search database with the standard PDB files from the PDB whose pocket structures were included in the PDBBind database. They were searched with foldseek2 program against the created database. Based on foldseek2 output, SMILES string information was retrieved per ligand listed in important sites (REMARK 800 in target PDB) when aligned portion between Alphafold structure of the homologous gene and target PDB included protein residues associated with the sites as listed in REMARK 800. IPD3 module information associated with the homologous gene was retrieved.

Secondary structure information and PDBBind flags of the residues were retrieved. True value was assigned to PDBBind flag if the aligned portion was overlapped with PDB pocket. We created a dataset where each instance included 476 variables of SMILES string, 1 variable of secondary structure information, 1 variable of IPD3 modules, and PDBBind flag (binary class). We excluded the instances where target chains were not A or where ligands were ions. WEKA software was used. Resample (supervised) filter and RandomForest classification algorithm were used [51]. Total number of instances was 191, and that of attributes was 479. The following default value parameter set was used: P (size of each bag: percent of training set size):100; I (number of iterations):100; Num-slots (number of execution slots):1; K (number of attributes to randomly investigate):0; M (minimum number of instances per leaf):1; V (minimum variance per split):0.001; S (seed for random number generator):1. 10 fold cross validation was used. Attribute selection algorithms in Weka (Best first, Gain Ratio feature evaluator, ReliefF Ranking Filter, Symmetrical Uncertainty Ranking Filter) with default values for parameters were used.

Protein structures of the histidine biosynthesis genes (AtHISN1B, AtHISN2, AtHISN1A, AtHISN6B, AtHISN3, AtHISN5A, AtHISN5B, AtHISN4, AtHISN6A, At-HISH, AtHISN8) were retrieved from Alphafold database, and searched against PDB structures belonging to the same and close COG numbers from M. truncatula. The same procedure as above was used for dataset creation except that no secondary structure variable was retrieved. We excluded the instances if their ligands were ions. The COG numbers of the query structures (A. thaliana) were COG0040, COG0079, COG0106, COG0107, COG0131, COG0139, COG0141, COG0311, while those of target structures (M. truncatula) were COG0040, COG0192, COG0345, COG0421, KOG3178, NOG040026. If the query and the target belonged to the same COG-COG0040, true value was assigned to the COG flag, the binary class. Total number of instances was 1020, and that of attributes was 478. The same algorithm, filter, and parameter set were used. Attribute selection algorithms (Best first, Correlation Ranking Filter, Information Gain Ranking Filter, ReliefF Ranking Filter) with default values for parameters were used.2.10.3 Analysis of DEGs from A. thaliana in AM network from Ivan Fernandez et al. [4]

We created functional groups: cytoskeleton, membrane transport, vesicular trafficking, secretion, subcellular localization sorting, primary/secondary/hormone metabolism, gene expression and RNA metabolism, protein synthesis and processing, signal transduction and post-translational regulation, defense and cell rescue, and abiotic stimuli and development [56]. From Go and PO files, we retrieved all genes whose descriptions included the words “polarity”, “phloem”, and “xylem” [53]. We retrieved PCSD chromatin state data, OCR sequence length, DDI category data (redundant and non-redundant DDI/DMI), max topology number of domain dimers, and pathway analysis data. We applied the clustering-canopy and cobweb with default value parameter sets in the Weka program.

From the string database, we added all genes that were predicted to have physical contact with DEGs. We created EWAS-phenotype data as described in the Muti-omics association module and applied the attribute selection method “CorrelationAttributeEval” in the Weka software. We excluded the variables containing only NA values and null values, namely: Enhancer_As75_CG, Enhancer_As75_CHG, Enhancer_As75_CHH, Enhancer_Cu65_CG, Enhancer_Cu65_CHG, Enhancer_Cu65_CHH, Enhancer_Ni60_CG, Enhancer_Ni60_CHG, Enhancer_Ni60_CHH, Enhancer_Mg25_CG, Enhancer_Mg25_CHG, Enhancer_Mg25_CHH, Enhancer_P31_CG, Enhancer_P31_CHG, and Enhancer_P31_CHH variables.

## 3 Results

The modules described in Methods section have been implemented in AraMultiOmics. Demonstration of the software usage, three simple analyses are shown below, where subsection numbers are paired with those of 2.10 in methods section.

### 3.1. Analysis of genes belonging to COGs with metals in their descriptions

Among COGs with descriptions containing at least one of four metals, we selected the families whose member genes distributed across various COG IDs: early nodulin-like protein, F-box protein, chaperone protein dnaJ, antiporter, calcium-binding, ATPase, ferredoxin, heavy metal-associated isoprenylated plant protein, peptidyl-prolyl cis-trans isomerase, probable WRKY TF, transmembrane protein, wall-associated receptor kinase, and auxin transporter. We further selected those with a large number of COGs: early nodulin-like protein, F-box protein, ATPase, and auxin transporter. COGs belonging to early nodulin-like protein and F-box protein families contained either up-regulated or down-regulated gene(s). In contrast, those belonging to ATPase and auxin transporters had no DEGs except COG0652. COG0652 contained mostly peptidyl-prolyl cis-trans isomerase CYP homologous genes, hydroxyproline-rich glycoprotein family protein, and leukocyte receptor cluster-like protein in addition to ATPase E1. IPD3 modules of the genes belonging to the same COGs varied widely in early nodulin-like protein and F-box protein. IPD3 modules in ATPase and auxin-related transport proteins were relatively restricted to a small number of IPD3 modules and contained no module S nor T. Transcription of these modules had a negative correlation with AMF treatment and transgenic Arabidopsis line expressing partial Medicago truncatula IPD3 (aa 254-513 of IPD3Mt). Associations between the DEG types of M. truncatula and IPD3 modules varied across gene families (supplementary data 1).

### 3.2. Analysis of homologous genes from A. thaliana with IPD3 restoration

Analysis with only PCSD chromatin state data had all homologous genes. FarthestFirst clustering method showed that all HAI, CYP76C, PAL, and CSLG homologous genes were grouped into one cluster, while two VTE homologous genes with IPD3 DEG modules B and E and two GUS homologous genes with D-A-B and E modules were grouped into the other cluster (supplementary data 2), which implies that chromatin states of VTE and GUS homologous genes may have different associations with AM colonization.

Clustering results applied to PCSD chromatin states and OCR SRA accession-tissue data in VOC pathways indicated that genes encoding proteins involved in the shikimate pathway belonged to one cluster, while PAL homologous genes belonged to different clusters, which implied that the distinct characteristics of the homologous genes were associated with epigenetic regulations and tissue type-specific properties.

Frequencies of DNA structures in PFK genes from two species showed minimal differences between the species (supplementary data 3). In both species, the frequencies of qquad (G-quadruplex) and slipped motifs located beyond 2000 bp were higher compared to other structures. Protein metal binding potentials of Ca, Cu, Fe, K, Mn, and Ni ions from M. truncatula varied widely compared to other ions (supplementary data 4).

Accuracy of the classification model created from the VOC dataset with 479 variables (SMILES string of the ligand, protein secondary structure, IPD3 modules) and the class of PDBBind flag was 96% (supplementary data 5). Kappa statistic was 0.84. Mean absolute error was 0.68. Model evaluation metrics were shown in the Table 1.

**Table 1.**
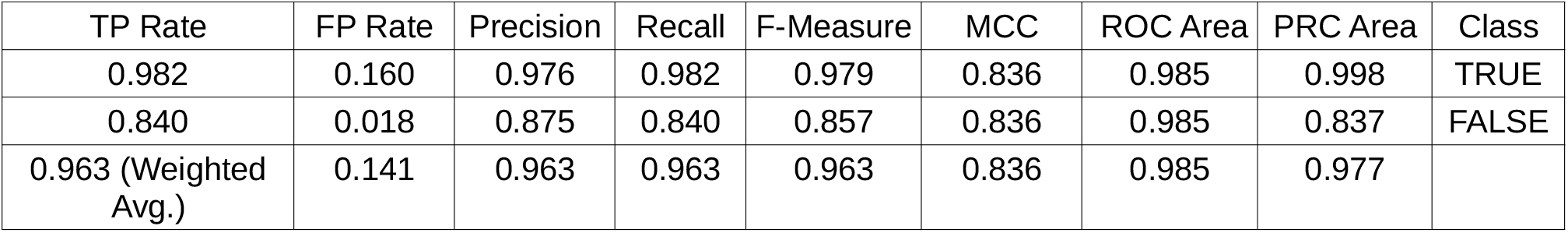
Model evaluation metrics for binary classification of the VOC dataset

Top 10 ranked attributes from four different attribute selection algorithms are shown in the Table 2.

**Table 2.**
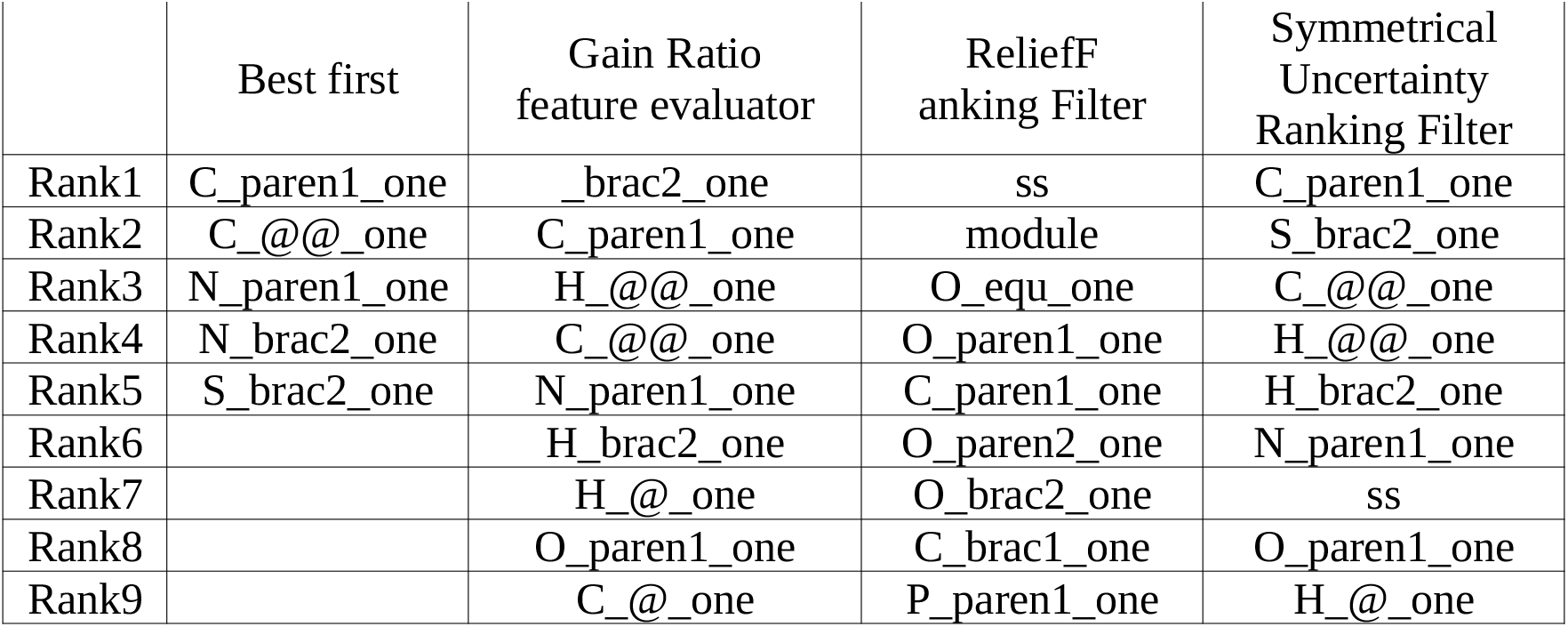
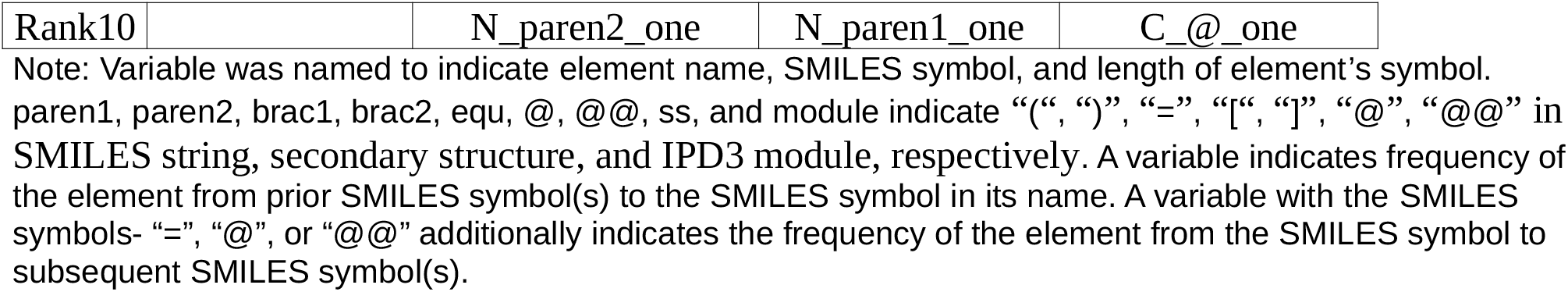
List of selected Top ranked attributes for PDBBind binary classification

|  | Best first | Gain Ratio<br>feature evaluator | ReliefF<br>anking Filter | Symmetrical<br>Uncertainty<br>Ranking Filter |
| --- | --- | --- | --- | --- |
| Rank1 | C_paren1_one | _brac2_one | ss | C_paren1_one |
| Rank2 | C_@@_one | C_paren1_one | module | S_brac2_one |
| Rank3 | N_paren1_one | H_@@_one | O_equ_one | C_@@_one |
| Rank4 | N_brac2_one | C_@@_one | O_paren1_one | H_@@_one |
| Rank5 | S_brac2_one | N_paren1_one | C_paren1_one | H_brac2_one |
| Rank6 |  | H_brac2_one | O_paren2_one | N_paren1_one |
| Rank7 |  | H_@_one | O_brac2_one | ss |
| Rank8 |  | O_paren1_one | C_brac1_one | O_paren1_one |
| Rank9 |  | C_@_one | P_paren1_one | H_@_one |
| Rank10 |  | N_paren2_one | N_paren1_one | C_@_one |

Note: Variable was named to indicate element name, SMILES symbol, and length of element's symbol. paren1, paren2, brac1, brac2, equ, @, @@, ss, and module indicate “(”, “)”, “=”, “[”, “]”, “@”, “@@” in SMILES string, secondary structure, and IPD3 module, respectively. A variable indicates frequency of the element from prior SMILES symbol(s) to the SMILES symbol in its name. A variable with the SMILES symbols- “=”, “@”, or “@@” additionally indicates the frequency of the element from the SMILES symbol to subsequent SMILES symbol(s).

Accuracy of the classification model created from the histidine biosynthesis gene dataset with 478 variables (SMILES string of the ligand, IPD3 modules) and the class of the same COG flag was 98% (supplementary data 6). Kappa statistic was 0.73. Mean absolute error was 0.037. Model evaluation metrics are shown in the Table 3.

**Table 3.**
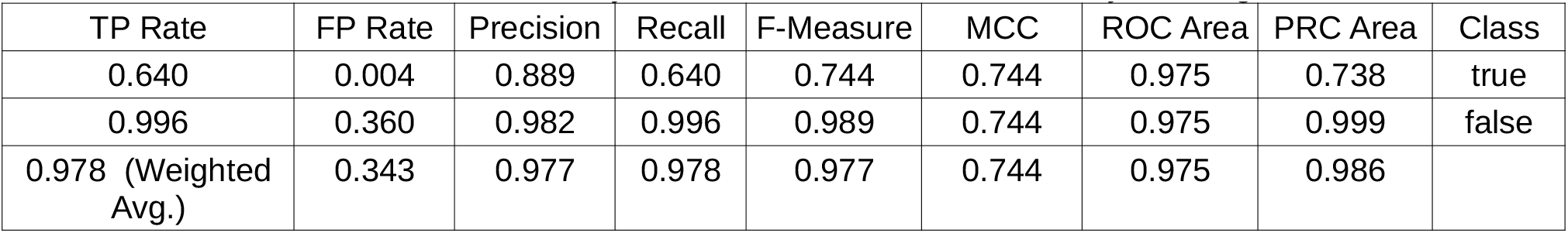
Model evaluation metrics for binary classification of the histidine biosynthesis gene dataset

When we removed IPD3 module variable, model performance significantly decreased. Top 10 ranked attributes from four different attribute selection algorithms are shown in the Table 4.

**Table 4.**
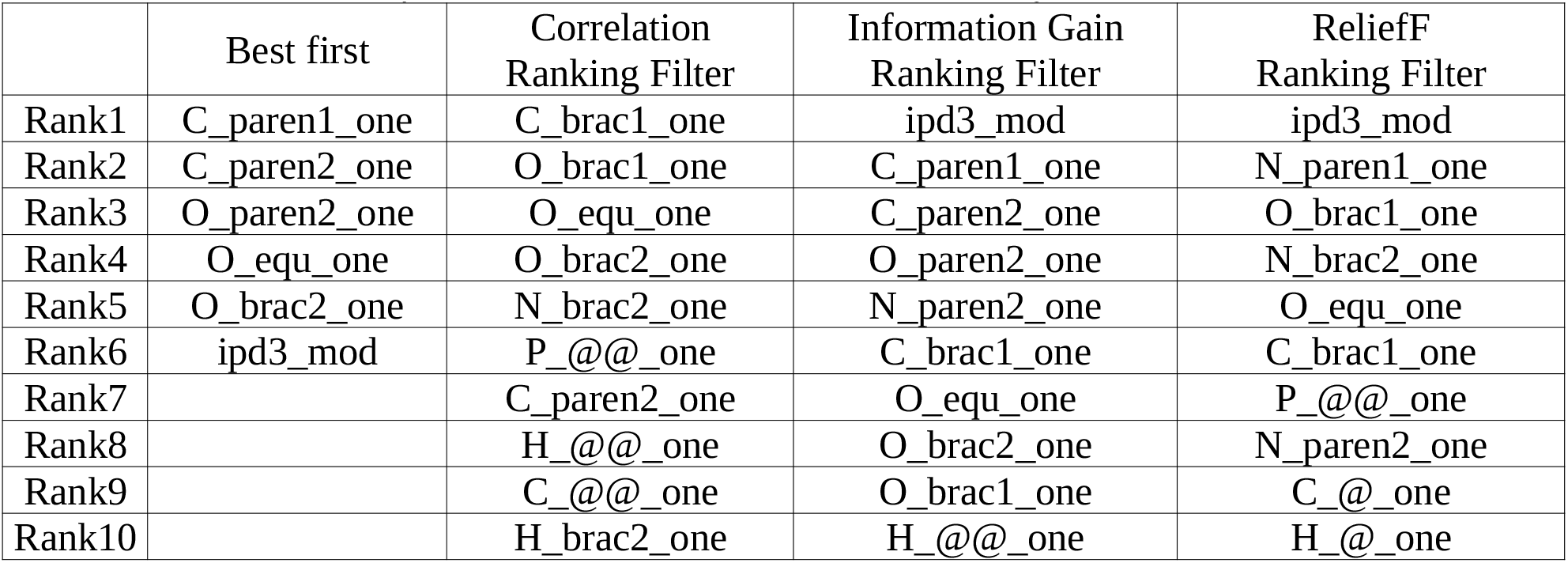
List of selected Top ranked attributes for the same COG binary classification

|  | Best first | Correlation Ranking Filter | Information Gain Ranking Filter | ReliefF Ranking Filter |
| --- | --- | --- | --- | --- |
| Rank1 | C_paren1_one | C_brac1_one | ipd3_mod | ipd3_mod |
| Rank2 | C_paren2_one | O_brac1_one | C_paren1_one | N_paren1_one |
| Rank3 | O_paren2_one | O_equ_one | C_paren2_one | O_brac1_one |
| Rank4 | O_equ_one | O_brac2_one | O_paren2_one | N_brac2_one |
| Rank5 | O_brac2_one | N_brac2_one | N_paren2_one | O_equ_one |
| Rank6 | ipd3_mod | P_@@_one | C_brac1_one | C_brac1_one |
| Rank7 |  | C_paren2_one | O_equ_one | P_@@_one |
| Rank8 |  | H_@@_one | O_brac2_one | N_paren2_one |
| Rank9 |  | C_@@_one | O_brac1_one | C_@_one |
| Rank10 |  | H_brac2_one | H_@@_one | H_@_one |

### 3.3. Analysis of DEGs from A. thaliana in AM network

We selected the PO numbers whose distributions in cluster memberships were distinct based on the cobweb clustering method applied to PCSD chromatin states and sequence length of accessible chromatin regions of the entire PO dataset. According to the PO descriptions, fractions of the distinct distribution in each group were calculated and sorted in descending order: secretion, vesicular, subcellular localization sorting, vacu, signal, phloem, defense, RNA, transport, transporter, metabolism, xylem (supplementary table 1,2, supplementary data 7,8).

Frequency table of group memberships in clusters, resulting from clustering DDI NR/non-NR category and of max topology number of domain dimer dataset, showed that all genes in the defense, metabolism, phloem, and xylem groups belonged to only one cluster, while those in the signal group distributed across all the clusters (supplementary table 3, supplementary data 9).

In the same vein, metabolic pathway analysis of DEG data indicated that the transporter group had diverse genes scattered throughout clusters while the xylum group had all members belonging to only one cluster (supplementary table 4, supplementary data 10).

Correlations of attributes of the EWAS-phenotype data of DEG from Ivan Fernandez et al. group showed that phenotypes-S34, Se82, and Mn55 were strongly correlated with each other, while As75, Co59, Cu65, Mo98, Ni60, P31, Zn66, Enhancer_Cd111_CHH, and Enhancer_Rb85_CHH were strongly correlated (supplementary table 5, supplementary data 11).

## 4 Discussions

AraMultiOmics program offers integrative resource for fast omics feature generation for downstream statistical analysis. Implementation of inter-module relational key-protein domain facilitates data retrieval and integration. PLP module needs upgrading for mapping pfam/interpro ID to PDB structures. Simple logic was used to generate variables from frequencies of atomic properties in SMILES string. We need to enhance the function to measure physicochemical properties of ligands with thermodynamic and kinetic data of protein-ligand interactions, a task that remains to be carried out in further research.

While data creation corresponding to PNI, multi-omics associations, and GO/PO/IPD3 is only available for the model species A. thaliana, the modules for DNA structure and metal binding profile, DNA-TF binding, DDI, and metabolic network analysis can be used for comparative study for multiple species. Comparative omics studies could help decipher the driving forces behind patterns of molecular interactions during AM symbiosis. Integration of PNI data for M. truncatular will be useful for direct comparison in gene regulation. For example, among many controlling mechanisms of arbuscule formation, the transcriptional negative feedback loop (ERM1/WRI5a-ERF12-TOPLESS) may fine-tune arbuscule development via PPI in M. truncatula [58]. With the integration of chip-seq, atac-seq, and TF binding site information for M. truncatula, the araMultiOmics and Art-foundation programs may provide valuable tools to assess the mechanisms underlying such transcriptional regulations [59]. The development of modules specific to M. truncatula remains a subject for further research.

Although DNA structures and protein metal binding profiles need to be analyzed on a genome-wide scale, sequence positions were not provided. In the same vein, the positions of secondary structures of membrane proteins were not supplied in protein orientation data.

Incorporation of the position information will improve the program’s performance. Unique features of AraMultiOmics include the integration of information of metal ion and ligand with complex structure, which can be coupled with various information, including epigenetic regulation, DNA structure, protein metal binding potential, gene expression, and metabolic pathway. It may enable researchers to make inferences for estimating the effects of fungal factors and root exudates of the AM network.

All the modules of AraMultiOmics were created based on existing databases and programs, some of which were mainly applied to mammals (e.g., DNA structure and metal binding profile, DNA-TF binding profiles, and Protein domain-domain interaction). Non-plant-specific databases and programs were integrated into AraMultiOmics due to the absence of such resources in the plant kingdom. Although integration of multiple omics data into the development of a prediction model may provide an effective means to infer the molecular basis of AM symbiosis of non-host plants, the interpretation should be taken with caution, considering the limitation of adopting a methodology for studying mammals to plant research. In a broader context, the issue of such limitation can be extended to existing molecular biology techniques and computational analysis approaches. Addressing these challenges requires the establishment of novel computational approaches that account for the inaccuracies arising from fundamental differences between mammals and plants. The development of computational methods to make proper adjustments remains a subject for further research.

## 6 Author Contributions

J.E.K.: conceptualization, methodology, software, formal analysis, writing-original draft, writing-review and editing.

## 8 Acknowledgments

I thank Mohamed Hijri for his guidance and support.

## 10 Supplementary Material

Supplementary Material are available online.

## 11 Data Availability Statement

The datasets generated and analyzed for this study can be found at www.artfoundation.kr.

**Fig 1.**
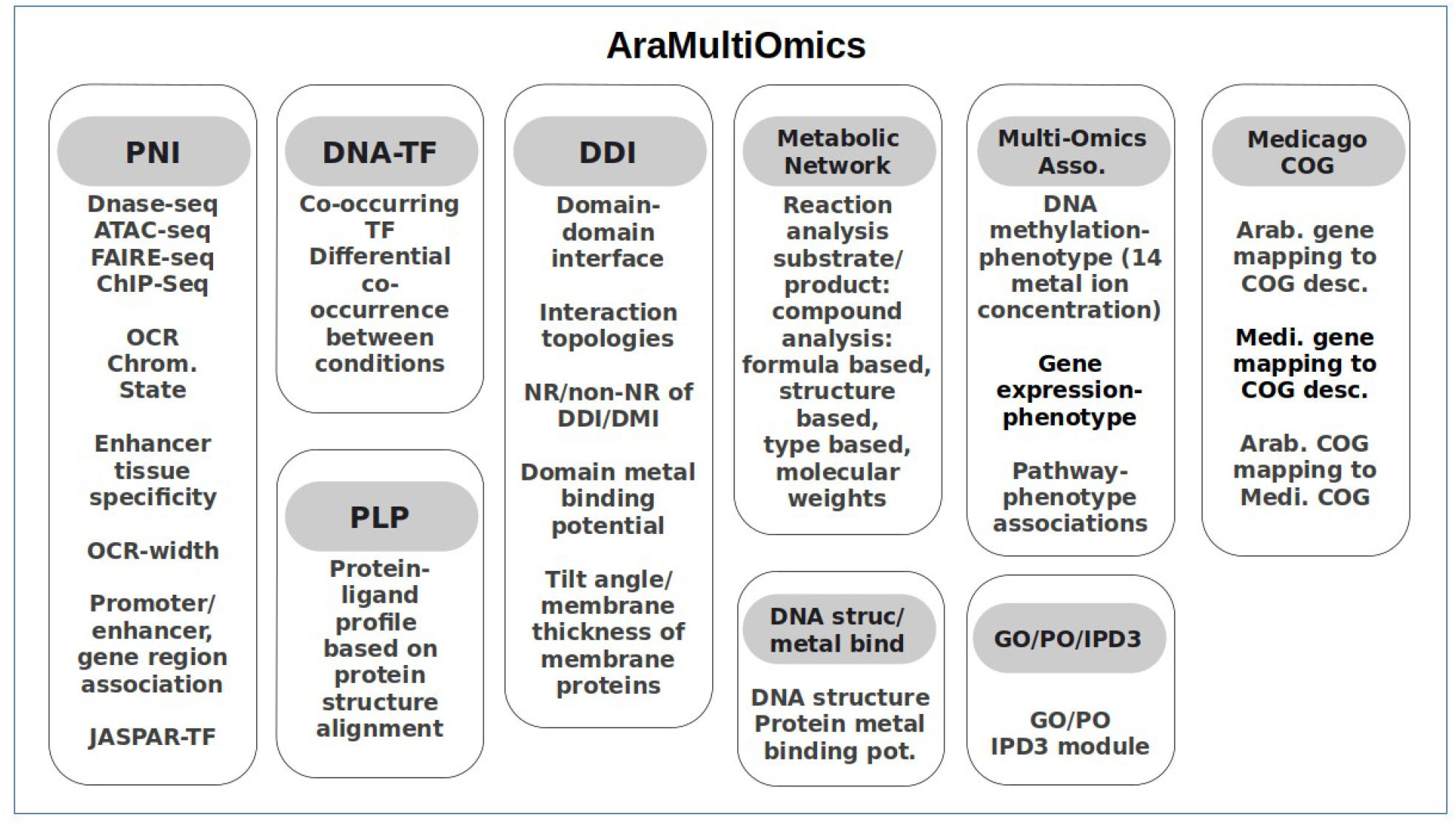
Program modules

## References

1. Kumar, V., & Kumar, P. (2019). A review on feasibility of phytoremediation technology for heavy metals removal. Archives of Agriculture and Environmental Science, 4(3), 326–341. 10.26832/24566632.2019.0403011

2. Yun HS, Kwaaitaal M, Kato N, Yi C, Park S, Sato MH, Schulze-Lefert P, Kwon C. (2013). Requirement of vesicle-associated membrane protein 721 and 722 for sustained growth during immune responses in Arabidopsis. Mol Cells. 2013 Jun;35(6):481–8. doi: 10.1007/s10059-013-2130-2. Epub 2013 May 8. PMID: 23661365; PMCID: PMC3887875.

3. Cosme M, Fernández I, Van der Heijden MGA, Pieterse CMJ. (2018). Non-Mycorrhizal Plants: The Exceptions that Prove the Rule. Trends Plant Sci. 2018 Jul;23(7):577–587. doi: 10.1016/j.tplants.2018.04.004. Epub 2018 May 9. PMID: 29753631.

4. Fernández, Iván, Cosme, Marco, Stringlis, Ioannis A., Yu, Ke, de Jonge, Ronnie, van Wees, SaskiaC.M., Pozo, Maria J., Pieterse, Corné M. J., van der Heijden, Marcel G. A. (2019). Molecular dialogue between arbuscular mycorrhizal fungi and the nonhost plant Arabidopsis thaliana switches from initial detection to antagonism. New Phytologist. 223.2. 0028–646X

5. Hornstein ED, Charles M, Franklin M, Edwards B, Vintila S, Kleiner M, Sederoff H. (2023). Re-engineering a lost trait: IPD3, a master regulator of arbuscular mycorrhizal symbiosis, affects genes for immunity and metabolism of non-host Arabidopsis when restored long after its evolutionary loss. bioRxiv [Preprint]. 2023 Mar 8:2023.03.06.531368. doi: 10.1101/2023.03.06.531368. PMID: 36945518; PMCID: PMC10028889.

6. Akihiro Yamazaki, Kai Battenberg, Yoshikazu Shimoda, and Makoto Hayashi. (2022). NDR1/HIN1-Like Protein 13 Interacts with Symbiotic Receptor Kinases and Regulates Nodulation in Lotus japonicus. MPMI Vol. 35, No. 9, pp. 845–856

7. Narusaka Y, Shinya T, Narusaka M, Motoyama N, Shimada H, Murakami K, Shibuya N. (2013). Presence of LYM2 dependent but CERK1 independent disease resistance in Arabidopsis. Plant Signal Behav. 2013 Sep;8(9):e25345. doi: 10.4161/psb.25345. Epub 2013 Aug 2. PMID: 23803749; PMCID: PMC4002583.

8. Quetglas S, Leveque C, Miquelis R, Sato K, Seagar M. (2000). Ca2+-dependent regulation of synaptic SNARE complex assembly via a calmodulin- and phospholipid-binding domain of synaptobrevin. Proc Natl Acad Sci U S A. 2000 Aug 15;97(17):9695–700. doi: 10.1073/pnas.97.17.9695. PMID: 10944231; PMCID: PMC16927.

9. Holland S, Roth R. (2023). Extracellular Vesicles in the Arbuscular Mycorrhizal Symbiosis: Current Understanding and Future Perspectives. Mol Plant Microbe Interact. 2023 Apr;36(4):235–244. doi: 10.1094/MPMI-09-22-0189-FI. Epub 2023 Apr 13. PMID: 36867731.

10. Liu Y, Xiong Z, Wu W, Ling HQ, Kong D. Iron in the Symbiosis of Plants and Microorganisms. Plants (Basel). 2023 May 11;12(10):1958. doi: 10.3390/plants12101958. PMID: 37653875; PMCID: PMC10223382.

11. Mélanie K. Rich, Eva Nouri, Pierre-Emmanuel Courty, Didier Reinhardt, Diet of Arbuscular Mycorrhizal Fungi: Bread and Butter?, Trends in Plant Science, Volume 22, Issue 8, 2017, Pages 652–660, ISSN 1360-1385, 10.1016/j.tplants.2017.05.008.

12. Meng Ying Gao, Xun Wen Chen, Wei Xiong Huang, Li Wu, Zheng Sheng Yu, Lei Xiang, Ce Hui Mo, Yan Wen Li, Quan Ying Cai, Ming Hung Wong, Hui Li, Cell wall modification induced by an arbuscular mycorrhizal fungus enhanced cadmium fixation in rice root, Journal of Hazardous Materials, Volume 416, 2021, 125894, ISSN 0304-3894, 10.1016/j.jhazmat.2021.125894.

13. Mikhail A. Lomize, Andrei L. Lomize, Irina D. Pogozheva, Henry I. Mosberg, OPM: Orientations of Proteins in Membranes database, Bioinformatics, Volume 22, Issue 5, March 2006, Pages 623–625, 10.1093/bioinformatics/btk023and

14. Fu, LY., Zhu, T., Zhou, X. et al. (2022). ChIP-Hub provides an integrative platform for exploring plant regulome. Nat Commun 13, 3413 10.1038/s41467-022-30770-1

15. Castro-Mondragon JA, Riudavets-Puig R, Rauluseviciute I, Berhanu Lemma R, Turchi L, Blanc-Mathieu R, Lucas J, Boddie P, Khan A, Manosalva Pérez N, Fornes O, Leung TY, Aguirre A, Hammal F, Schmelter D, Baranasic D, Ballester B, Sandelin A, Lenhard B, Vandepoele K, Wasserman WW, Parcy F, and Mathelier A JASPAR 2022: the 9th release of the open-access database of transcription factor binding profiles Nucleic Acids Res. journal_details; doi: 10.1093/nar/gkab1113

16. Liu Y, Tian T, Zhang K, You Q, Yan H, Zhao N, Yi X, Xu W, Su Z. (2018). PCSD: a plant chromatin state database. Nucleic Acids Res. 2018 Jan 4;46(D1):D1157–D1167. doi: 10.1093/nar/gkx919. PMID: 29040761; PMCID: PMC5753246.

17. Tan G, Lenhard B. (2016). TFBSTools: an R/bioconductor package for transcription factor binding site analysis. Bioinformatics. 2016 May 15;32(10):1555–6. doi: 10.1093/bioinformatics/btw024. Epub 2016 Jan 21. PMID: 26794315; PMCID: PMC4866524.

18. Higo, Kenichi, Ugawa, Yoshihiro, Iwamoto, Masao, Higo, Hiromi. (1998). PLACE: A database of plant cis -acting regulatory DNA elements. Nucleic Acids Research. 1998. 26. 1. 358–9. 10.1093/nar/26.1.358

19. Kikin O, D’Antonio L, Bagga PS. (2006). QGRS Mapper: a web-based server for predicting G-quadruplexes in nucleotide sequences. Nucleic Acids Res. 2006 Jul 1;34(Web Server issue):W676–82. doi: 10.1093/nar/gkl253. PMID: 16845096; PMCID: PMC1538864.

20. Aptekmann AA, Buongiorno J, Giovannelli D, Glamoclija M, Ferreiro DU, Bromberg Y. (2022). mebipred: identifying metal-binding potential in protein sequence. Bioinformatics. 2022 Jul 11;38(14):3532–3540. doi: 10.1093/bioinformatics/btac358. PMID: 35639953; PMCID: PMC9272798.

21. Mette Bentsen, Vanessa Heger, Hendrik Schultheis, Carsten Kuenne, Mario Looso, (2022). TF-COMB – Discovering grammar of transcription factor binding sites, Computational and Structural Biotechnology Journal, Volume 20, 2022, Pages 4040–4051, ISSN 2001-0370

22. Mosca, R.; Céol, A.; Stein, A.; Olivella, R.; Aloy, P. (2013). 3did: A catalog of domain-based interactions of known three-dimensional structure. Nucleic Acids Res. 2013, 42, D374–D379, 10.1093/nar/gkt887.

23. Jumper, J et al. Highly accurate protein structure prediction with AlphaFold. Nature (2021). Varadi, M et al. AlphaFold Protein Structure Database: massively expanding the structural coverage of protein-sequence space with high-accuracy models. Nucleic Acids Research (2021).

24. van Kempen M, Kim S, Tumescheit C, Mirdita M, Lee J, Gilchrist CLM, Söding J, and Steinegger M. Fast and accurate [protein structure search with Foldseek. Nature Biotechnology, 2023.

25. H.M. Berman, J. Westbrook, Z. Feng, G. Gilliland, T.N. Bhat, H. Weissig, I.N. Shindyalov, P.E. Bourne, The Protein Data Bank (2000) Nucleic Acids Research 28: 235–242 10.1093/nar/28.1.235.

26. Wang, R.; Fang, X.; Lu, Y.; Wang, S. “The PDBbind Database: Collection of Binding Affinities for Protein-Ligand Complexes with Known Three-Dimensional Structures”, J. Med. Chem., 2004; 47(12); 2977–2980. (PDBbind prototype)

27. Delphine Bahammou, Ghislaine Recorbet, Adiilah Mamode Cassim, Franck Robert, Thierry Balliau, Pierre Van Delft et al. Revealing the lipidome and proteome of Arabidopsis thaliana plasma membrane. BioRxiv 2023.05.14.540643; doi: 10.1101/2023.05.14.540643

28. Yamada I, Shiota M, Shinmachi D, Ono T, Tsuchiya S, Hosoda M, Fujita A, Aoki NP, Watanabe Y, Fujita N, Angata K, Kaji H, Narimatsu H, Okuda S, Aoki-Kinoshita KF. The GlyCosmos Portal: a unified and comprehensive web resource for the glycosciences. Nat Methods. 2020 Jul;17(7):649–650. doi: 10.1038/s41592-020-0879-8. PMID: 32572234.

29. Yanbin Yin, Xizeng Mao, Jincai Yang, Xin Chen, Fenglou Mao, Ying Xu, dbCAN: a web resource for automated carbohydrate-active enzyme annotation, Nucleic Acids Research, Volume 40, Issue W1, 1 July 2012, Pages W445–W451, 10.1093/nar/gks479

30. Finn RD, Clements J, Eddy SR. HMMER web server: interactive sequence similarity searching. Nucleic Acids Res. 2011 Jul;39(Web Server issue):W29–37. doi: 10.1093/nar/gkr367. Epub 2011 May 18. PMID: 21593126; PMCID: PMC3125773.

31. Hawkins, C., Ginzburg, D., Zhao, K., Dwyer, W., Xue, B., Xu, A., Rice, S., Cole, B., Paley, S., Karp, P., Rhee, S. Y. (2021). Plant Metabolic Network 15: A resource of genome-wide metabolism databases for 126 plants and algae. Journal of Integrative Plant Biology. [HTML]

32. Lan, Yiheng, Sun, Ruikun, Ouyang, Jian, Ding, Wubing, Kim, Min-Jun, Wu, Jun, Li, Yuhua, Shi, Tieliu. (2020). AtMAD: Arabidopsis thaliana multi-omics association database. Nucleic Acids Research. 2020. 49. D1. D1445–D1451. 10.1093/nar/gkaa1042

33. Tian H, Li Y, Wang C, Xu X, Zhang Y, Zeb Q, Zicola J, Fu Y, Turck F, Li L, Lu Z, Liu L. (2021). Photoperiod-responsive changes in chromatin accessibility in phloem companion and epidermis cells of Arabidopsis leaves. Plant Cell. 2021 May 5;33(3):475–491. doi: 10.1093/plcell/koaa043. PMID: 33955490; PMCID: PMC8136901.

34. Seren Ü, Grimm D, Fitz J, Weigel D, Nordborg M, Borgwardt K, Korte A. AraPheno: a public database for Arabidopsis thaliana phenotypes. Nucleic Acids Res. 2017 Jan 4;45(D1):D1054–D1059. doi: 10.1093/nar/gkw986. Epub 2016 Oct 24. PMID: 27924043; PMCID: PMC5210660.

35. Berardini, T.Z.; Reiser, L.; Li, D.; Mezheritsky, Y.; Muller, R.; Strait, E.; Huala, E. The arabidopsis information resource: Making and mining the “gold standard” annotated reference plant genome. Genesis 2015, 53, 474–485, 10.1002/dvg.22877.

36. Szklarczyk, D.; Gable, A.L.; Lyon, D.; Junge, A.; Wyder, S.; Huerta-Cepas, J.; Simonovic, M.; Doncheva, N.T.; Morris, J.H.; Bork, P.; et al. STRING v11: Protein–protein association networks with increased coverage, supporting functional discovery in genome-wide experimental datasets. Nucleic Acids Res. 2019, 47, D607–D613, 10.1093/nar/gky1131.

37. R Core Team (2022). R: A language and environment for statistical computing. R Foundation for Statistical Computing, Vienna, Austria. URL https://www.R-project.org/.

38. Bashir K, Rasheed S, Kobayashi T, Seki M, Nishizawa NK. Regulating Subcellular Metal Homeostasis: The Key to Crop Improvement. Front Plant Sci. 2016 Aug 5;7:1192. doi: 10.3389/fpls.2016.01192. PMID: 27547212; PMCID: PMC4974246.

39. Radrich, K., Tsuruoka, Y., Dobson, P. et al. Integration of metabolic databases for the reconstruction of genome-scale metabolic networks. BMC Syst Biol 4, 114 (2010). 10.1186/1752-0509-4-114

40. Burks DJ, Sengupta S, De R, Mittler R, Azad RK. The Arabidopsis gene co-expression network. Plant Direct. 2022 Apr 26;6(4):e396. doi: 10.1002/pld3.396. PMID: 35492683; PMCID: PMC9039629.

41. Egorova KS, Smirnova NS, Toukach PV. CSDB_GT, a curated glycosyltransferase database with close-to-full coverage on three most studied nonanimal species. Glycobiology. 2021 Jun 3;31(5):524–529. doi: 10.1093/glycob/cwaa107. PMID: 33242091.

42. Ashburner, M., Ball, C., Blake, J. et al. Gene Ontology: tool for the unification of biology. Nat Genet 25, 25–29 (2000). 10.1038/75556.

43. Wu QS, Silva FSB, Hijri M, Kapoor R (2023) Editorial: Arbuscular mycorrhiza-mediated augmentation of plant secondary metabolite production. Front Plant Sci. 14:1150900. doi: 10.3389/fpls.2023.1150900.

44. Blakney AJC, Bainard LD, St-Arnaud M, Hijri M (2022) Soil Chemistry and Soil History Significantly Structure Oomycete Communities in Brassicaceae Crop Rotations. Applied and Environmental Microbiology, 89 (1)

45. Hijri M (2023) Microbial-Based Plant Biostimulants. Microorganisms 11(3):686.

46. Zambelli B, Musiani F, Ciurli S. Metal ion-mediated DNA-protein interactions. Met Ions Life Sci. 2012;10:135–70. doi: 10.1007/978-94-007-2172-2_5. PMID: 22210338.

47. Song WJ, Sontz PA, Ambroggio XI, Tezcan FA. Metals in protein-protein interfaces. Annu Rev Biophys. 2014;43:409–31. doi: 10.1146/annurev-biophys-051013-023038. PMID: 24773016.

48. Zhu H, Kim DJ, Baek JM, Choi HK, Ellis LC, Küester H, McCombie WR, Peng HM, Cook DR. Syntenic relationships between Medicago truncatula and Arabidopsis reveal extensive divergence of genome organization. Plant Physiol. 2003 Mar;131(3):1018–26. doi: 10.1104/pp.102.016436. PMID: 12644654; PMCID: PMC166867.

49. RCLASS: Reaction Classification and Ontology. Bioinformatics Center, Institute for Chemical Research, Kyoto University, Uji, Kyoto 611-0011, Japan. Data accessed on Oct. 20, 2023. https://www.genome.jp/reaction/

50. Y. Sulaiman, Bin Liu, Eve Kaurilind, Ülo Niinemets. Phloem-feeding insect infestation antagonizes volatile organic compound emissions and enhances heat stress recovery of photosynthesis in Origanum vulgare. Environmental and Experimental Botany. Volume 189, September 2021, 104551

51. Frank, E.; Hall, M.A.; Witten, I.H. The WEKA Workbench. In Data Mining: Practical Machine Learning Tools and Techniques, 4th ed.; Morgan Kaufmann: Burlington, MA, USA, 2016.

52. D. Fisher (1987). Knowledge acquisition via incremental conceptual clustering. Machine Learning. 2(2):139–172.

53. J. H. Gennari, P. Langley, D. Fisher (1990). Models of incremental concept formation. Artificial Intelligence. 40:11–61.

54. A. McCallum, K. Nigam, L.H. Ungar: Efficient Clustering of High Dimensional Data Sets with Application to Reference Matching. In: Proceedings of the sixth ACM SIGKDD internation conference on knowledge discovery and data mining ACM-SIAM symposium on Discrete algorithms, 169–178, 2000.

55. Leo Breiman (2001). Random Forests. Machine Learning. 45(1):5–32.

56. Hohnjec N, Czaja-Hasse LF, Hogekamp C, Küster H. Pre-announcement of symbiotic guests: transcriptional reprogramming by mycorrhizal lipochitooligosaccharides shows a strict co-dependency on the GRAS transcription factors NSP1 and RAM1. BMC Genomics. 2015 Nov 23;16:994. doi: 10.1186/s12864-015-2224-7. PMID: 26597293; PMCID: PMC4657205.

57. Fisher K, Turner S. PXY, a receptor-like kinase essential for maintaining polarity during plant vascular-tissue development. Curr Biol. 2007 Jun 19;17(12):1061–6. doi: 10.1016/j.cub.2007.05.049. PMID: 17570668.

58. Zhang, Q., Wang, S., Xie, Q. et al. Control of arbuscule development by a transcriptional negative feedback loop in Medicago. Nat Commun 14, 5743 (2023). 10.1038/s41467-023-41493-2

59. Kang JE, Jun JH, Kwon JH, Lee JH, Hwang K, Kim S, Jeong N. Arabidopsis Transcription Regulatory Factor Domain/Domain Interaction Analysis Tool-Liquid/Liquid Phase Separation, Oligomerization, GO Analysis: A Toolkit for Interaction Data-Based Domain Analysis. Genes (Basel). 2023 Jul 19;14(7):1476. doi: 10.3390/genes14071476. PMID: 37510380; PMCID: PMC10379056.

